# Vascular Biomechanics and Brain Biochemistry in Aged and Alzheimer’s Disease Mouse Models

**DOI:** 10.1101/2025.03.27.645832

**Authors:** Allison R. Jones, Amin Jarrahi, Kylee Karpowich, Lindsay Brown, Kalynn M. Schulz, Rebecca A. Prosser, A. Colleen Crouch

## Abstract

Age-related vascular changes accompany or precede the development of Alzheimer’s disease (AD) pathology. The comorbidity of AD and arterial stiffening may suggest that vascular changes have a pathogenic role. Carotid artery mechanics and hemodynamics have been associated with age-related cognitive decline. However, the impact of hemodynamics and vascular mechanics on regional vulnerability within the brain have not been thoroughly explored. Despite the venous system’s role in transport, the impact of age-related alterations of the brain venous circulation on cognitive impairment is much less understood compared to the arterial system. By studying vascular mechanics and the resulting spatially-resolved brain lipids in young and aged AD mice, we can determine the relationship between vascular stiffening and brain function. Young and aged female 3xTg mice and age-matched controls were imaged using a combination of ultrasound and mass spectrometry. Wall shear stress varied across age and AD (p<0.05). The circumferential cyclic strain values for the carotid arteries and the WSS values for the jugular veins between groups were measured but were not statistically significant. Both mean velocity and pulsatility index (PI) varied across age and AD (p<0.05). Liquid chromatography mass spectrometry (LC-MS) of brain tissue identified several lipids and metabolites with statistically significant quantities (p<0.05). The fold change was computed for young AD vs. young control, aged AD vs. aged control, aged control vs. young control, and aged AD vs. young AD. The abundance of several lipid headgroups changed significantly with respect to age and AD. Phosphatidylcholines (PC), phosphatidylethanolamines (PE), cardiolipins (CL), phosphatidylserines (PS), and lysophosphatidylcholines (LPC) have been shown to decrease with AD in previous studies. However, we observed a statistically significant increase in PC, PE, CL, PS, and LPC in the aged 3xTG mouse model compared to aged controls. Hexosylceramides (HexCer), ceramides (CER) and sphingomyelin (SM), classes of sphingolipids; lysophosphatidylethanolamine (LPE), a class of phospholipids; and onogalactosyl diglycerides (MGDG), a class of glycerolipids, have been shown to increase with AD in previous studies which aligns with the statistically significant increase of LPC, HexCer, CER, SM, LPE, and MDG observed in the aged 3xTg group compared to controls in this work. Combining both ultrasound imaging and mass spectrometry, we were able determine significant differences in the vascular biomechanics and brain biochemistry seen with aging and AD.

## Introduction

In 2024, approximately 7 million people in the United States were living with Alzheimer’s disease, and by 2060 this number is projected to nearly triple to 14 million [1][2]. Alzheimer’s disease (AD) is the most common type of dementia and is responsible for progressive memory loss, cognitive dysfunction, and behavioral changes [3]. AD is currently a leading cause of death worldwide in part due to the lack of effective treatments [4]. AD has been characterized by an accumulation of amyloid plaques and neurofibrillary tangles in the brain [4]; however, current medications have not been shown to prevent onset or reverse the progression of cognitive decline. Currently, the only treatment available is designed to slow the progression of the disease by targeting amyloid plaques [5][6]. To develop a treatment for AD, the causes of the disease must be thoroughly understood. One promising avenue of research is the connection between cardiovascular function and neurological disorders. In 2019, a meta-analysis showed that increased arterial stiffness is associated with markers of cerebral small-vessel disease and cognitive decline [7].

The comorbidity of AD and pathologies such as arterial stiffening may suggest that vascular changes have a pathogenic role [5][8][4]. Carotid artery mechanics (increased stiffness[5]) and hemodynamics (lower wall shear stress[9]) have been associated with age-related cognitive decline. However, the impact of hemodynamics and vascular mechanics on regional vulnerability within the brain have not been thoroughly explored. Downstream vascular effects, including increased jugular venous reflux, are associated with white matter disease and aging [10]. Despite the venous system’s role in transport, the impact of age-related alterations of brain venous circulation on cognitive impairment and dementia is much less understood compared to the arterial system [11]. Additionally, vascular changes accompany or precede the development of AD’s pathology and could be used as a biomarker for age-related dementia [12].

Lipids are crucial for a wide range of biological functions, including energy storage, cellular structure, and signaling processes. Due to their involvement in these essential physiological activities, the field of lipidomics has developed to study and analyze lipid-related processes. The molecular changes that lead to arterial stiffening and AD have been independently studied. Lipid alterations, in particular, have been linked to both arterial stiffening and AD [13], [14], [15], [16], [17], [18] [19][20], [21]. Not only are lipids relevant in vascular dysfunction, but lipid dysregulation and composition are implicated in various neurological disorders [13] [22][23] [24]. The brain is one of the most lipid-rich organs, which makes understanding the lipids involved in brain-related disease such as AD vital for understanding the disease itself.

To understand vascular implications on brain molecules such as lipids, mouse models genetically modified to develop AD are useful because of their accelerated onset of disease pathology and are currently used by many research groups to study AD [25][26], [27]. One of the most popular mouse models the 3xTg-AD transgenic model that expresses a human tauopathy mutation (tauP301L) in addition to APP (K670N/M671L) and PS1 (M146V) mutations associated with amyloid pathology (Jackson Laboratory) [25], [28], [29]. Only female mice were used in this study because of the decrease in phenotypic expression in male mice reported to Jackson Laboratories in 2014 by the donating investigator. Since women are more at risk of developing AD, with two-thirds of patients with AD being women, studying a female AD model will provide more insight into the disease [1][31].

In this work, we used ultrasound to assess the biomechanics of the vasculature, MALDI MSI to spatially-resolve lipid changes in the brain, and LC-MS to annotate the lipids by measuring their retention time. By combining ultrasound, MALDI mass spectrometry imaging, LC-MS, and histology data, we can understand how aging and AD affect the relationship between the vasculature and the brain. With the growth of an aging population and the increasing burden of health care for individuals with AD, studying the causes of AD is vital to reduce morbidity and mortality [32].

## Materials and Methods

All animal work was approved by the Institutional Animal Care and Use Committee (IACUC). Young (12 weeks) and aged (52 weeks) female B6;129-Tg(APPSwe,tauP301L)1Lfa *Psen1^tm1Mpm^* (3xTg) mice and age-matched controls (Jackson Laboratory) were used for this experiment [28]. Female 3xTg mice were used because the males have been shown not to develop the phenotypic traits of Alzheimer’s disease (AD) [29]. The average weights are reported in Table 1 (p<0.0001). An overview of the methods is depicted in Figure 1. The vasculature of the mice was imaged via ultrasound (Vevo 3100, Visual Sonics). After conscious decapitation, the brains were removed for analysis with mass spectrometry.

**Figure 1.**
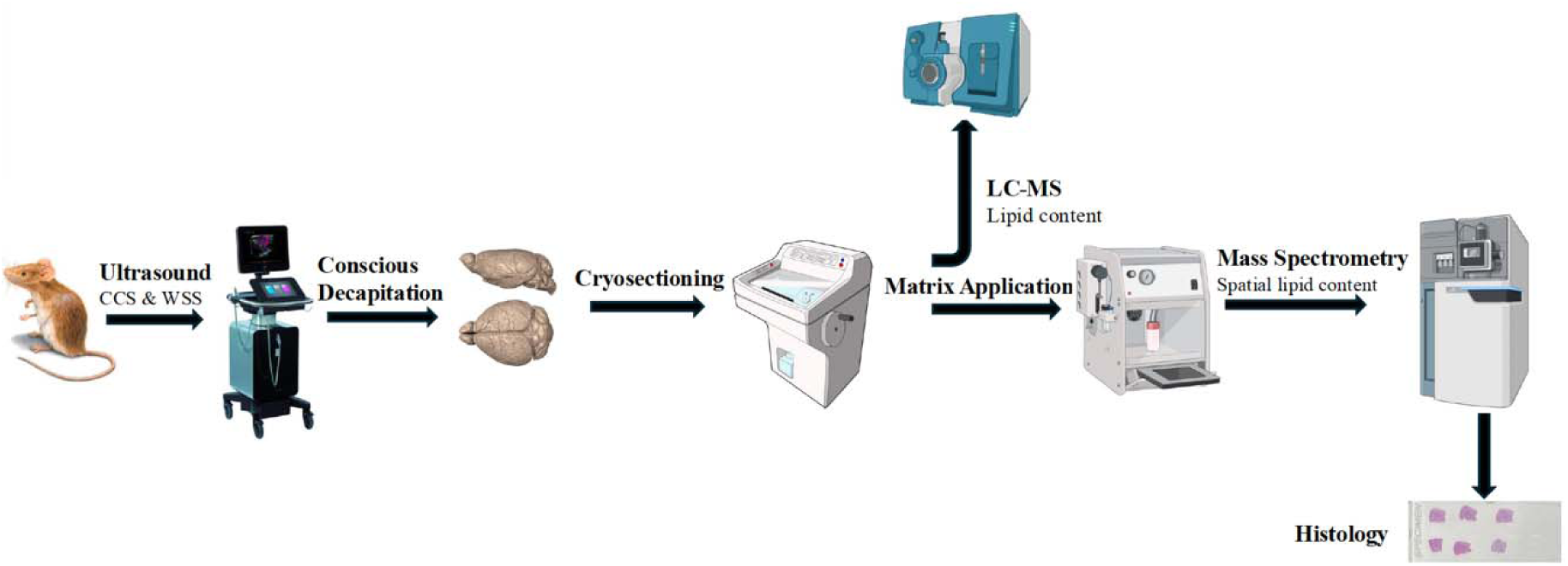

**Table 1:** Mouse Body Mass.

| Mouse | Average Mass (g) |
| --- | --- |
| YFM C*$\diamond$ | 23.0 ± 1.41 |
| OFM C*$\nabla\alpha$ | 45.2 ± 5.23 |
| YFM AD$^{\nabla}$ | 22.8 ± 2.14 |
| OFM AD$^{\diamond\alpha}$ | 53.2 ± 7.63 |

The aged 3xTg and aged control were statistically significant with $p = 0.0417$ . Symbols represent statistically significant differences between pairs.

### Ultrasound

Mice were imaged via ultrasound using the Vevo 3100 with the MX550D: 55 MHz MX Series transducer and analyzed with VevoLab (Visual Sonics). All images were collected with a gain setting of 35 dB. Images were collected in B-mode to measure the diameter of the carotid arteries and jugular veins (Figure 2). The diameter of the carotid artery was measured in three different locations directly inferior to the carotid bifurcation. The average diameter of the artery was calculated for both systole and diastole. The diameter of the jugular vein was also measured at three different time points across the cardiac cycle to provide an average diameter. The values of the diameters were used to calculate the circumferential cyclic strain (E) using the following equation,

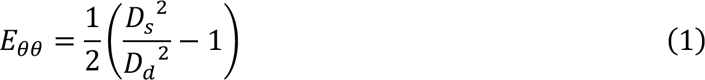

where *D_s_* represents the diameter during systole and *D_d_* represents the diameter during diastole for circular vessels [33] [34] [35]. The blood velocity through the carotid and jugular was measured using pulsed wave (PW) Doppler mode during systole and diastole (Figure 2). For the carotid artery, three velocity peaks in systole across the cardiac cycle were measured and averaged to provide an average velocity and the process was repeated in diastole. For the jugular vein, three velocity measurements were averaged to provide an average velocity. Using the velocity and diameter for the carotid in systole and diastole, the wall shear stress (WSS) was calculated. WSS was calculated using the following equation,

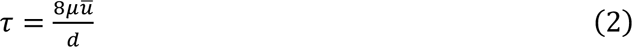

where µ is the viscosity of blood, 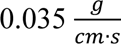, *u* is the average velocity of blood, and *d* is the average diameter [36] [37] [38][39]. In previous studies, the Hagen-Poiseuille application [40] has been used for the WSS measurements in large arteries. Since large arteries, such as the carotid artery, mimic the structure of a cylindrical tube with rigid walls, this technique is an appropriate method due to its simplicity [36][41]. WSS was not measured in the jugular veins because veins are generally a low-pressure, low-flow vessel compared to the arteries, do not mimic cylindrical tubes, and do not typically pulsate in the same way [42]. For the carotid arteries, the mean velocity was measured and used to calculate the PI using the following equation,

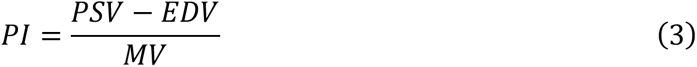

Where PSV is the peak systolic velocity, EDV is the end diastolic velocity, and MV is the mean velocity across the cardiac cycle [43]. PI is a non-invasive method of measuring vascular resistance with higher values indicating greater pulsatile flow possibly indicating vascular stiffening. [44]

**Figure 2.**
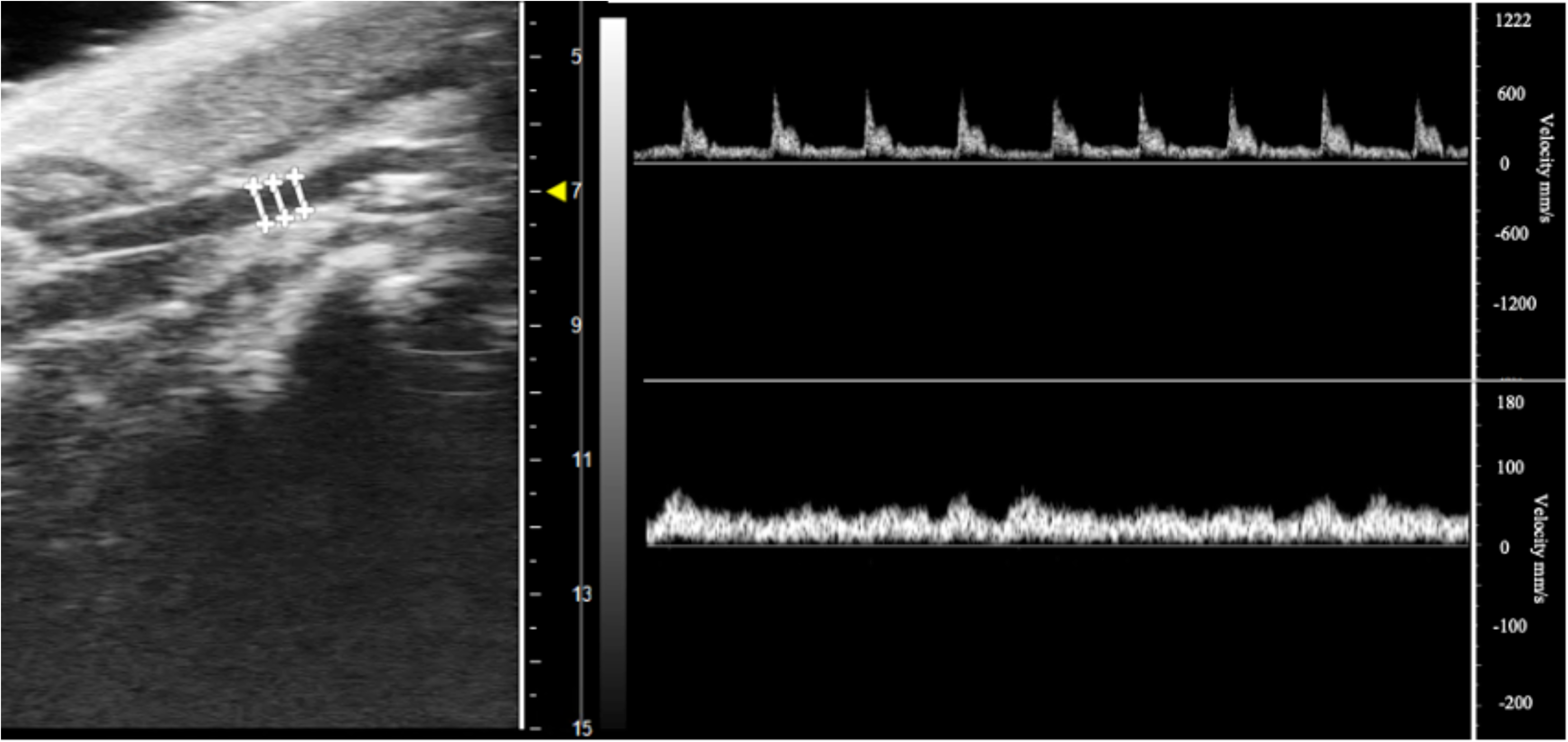
B-mode image (left) and pulse waves of the carotid artery (top right) and jugular vein (bottom right) of a young 3xTg control mouse with diameter measurements shown. The three values of diameter were averaged to obtain the average diameter of the carotid artery and used to calculate the CCS and WSS.

#### Statistical Analysis

Using JMP, a one-way ANOVA and a Tukey’s Honest Significance Difference (HSD) post-hoc test was performed to determine statistical difference between age and AD versus control groups for circumferential cyclic strain (CCS), WSS in systole and diastole for the carotid, and WSS for the jugular vein. Data is reported as mean and standard error for all data sets. Significance was set to p< 0.05.

### Tissue Harvesting and Preparation

After ultrasound, mice were sacrificed via conscious decapitation for quick removal of the brain. Conscious decapitation does not require the use of an anesthetic such as isoflurane, which can have an effect on the brain biochemistry [45]. The brains were cut in half and the left hemisphere was used by another research group. The right hemisphere of the brains was then flash frozen on liquid nitrogen by floating the tissue in aluminum foil boats. The brains were cryosectioned at 10 *μ*m and mounted on positively charged glass microscope slides for MALDI MSI and serial sections (∼60 *μ*m) were collected in tubes for LC-MS analysis. The slides and tubes were stored in a -80° C freezer until used for mass spectrometry.

### Mass Spectrometry

#### MALDI-MSI

2,5-Dihydroxybenzoic acid (DHB) was sprayed on the slides using an HTX M3+ sprayer, with spray parameters listed in Table 2. The matrix was sprayed on the tissue in a crisscross pattern 10 times at a temperature of 75° C and a pressure of 10 psi N2 gas. Once the tissue was sprayed with the matrix, the slide was scanned (Epson Scanner) and then was placed in a Waters Synapt mass spectrometer and run in positive mode with a pixel size set to 60 μm, 300 laser shots fired at 1kHz using a laser pulse energy with an average of 25 μJ. The laser was adjusted to 300 arbitrary units (a.u.) [46]. The mass spectra were processed using MassLynx software and the resulting molecular ion images were analyzed using HD Imaging (Waters Corp).

**Table 2:** Spray parameters for MALDI MSI using M3+ HTX Sprayer.

| **Matrix** | **Polarity** | **Solvent** | **Conc.**<br>(mg/mL) | **Flow**<br>**Rate**<br>($\mu\text{m}/\text{min}$) | **Velocity**<br>(mm/min) | **Track**<br>**Spacing**<br>(mm) | **Drying**<br>**Time**<br>(s) |
| --- | --- | --- | --- | --- | --- | --- | --- |
| DHB | Positive | 70%<br>MeOH | 40 | 100 | 1200 | 3 | 10 |

#### LC-MS

LC-MS was performed by the Biological and Small Molecule Mass Spectrometry Core (BSMMSC) of The University of Tennessee Knoxville to identify lipids and metabolites in the brain samples. The brain samples were first homogenized followed by a protocol for lipid extraction. The protocol consisted of a liquid-liquid extraction with 2:1 ratio of methanol chloroform to water. The extracts were then dried under N2 and resuspended in 9:1 ratio of methanol to chloroform. The extracted lipids were separated using a Thermo Accucore C30 column on a Vanquish UHPLC coupled to an Exploris 120 Mass Spectrometer. The LC-MS data provides retention time allowing for more accurate lipid identification in comparison to MALDI MSI [47]. The lipids identified from the LC-MS analysis were compared with those found from the MALDI MSI analysis to search for similarities by importing the list of m/z values into HD imaging and reprocessing the raw data. The lipids that were found in both the LC-MS and MALDI MSI data and in all six mice for each group were analyzed further and compared to the results of previous studies.

### Statistical Analysis

Mass spectrometry statistics were calculated using MetaboAnalyst. Principal component analysis (PCA) is an unsupervised dimensionality reduction technique that was used to determine variability in lipid intensity between the groups [48]. Clustering was used to group similar lipid profiles together to identify patterns amongst the different groups [49]. Partial least squares discriminant analysis (PLS-DA) is a supervised dimensionality reduction technique that was used to separate all four groups[50].

### Staining

After MALDI-MSI, the slides were placed in a desiccator prior to staining, as seen in Figure 1. First, the matrix was removed using ethanol and HPLC water. After removing the matrix, the slide was air-dried for 5 minutes before beginning the staining protocol. The brain was stained using the Visium H&E staining protocol [51]. The slide was coated with isopropanol and left to sit for 1 minute. Isopropanol was removed and the slide was coated in hematoxylin and left to sit for 7 minutes. The hematoxylin was removed, and the slide was submerged in HPLC water. The slide was then coated in bluing buffer for 2 minutes and then submerged in HPLC water. Eosin mix was prepared using a combination of tris acetic acid and Eosin Y. The slide was coated in the eosin mix for 1 minute and then submerged in HPLC water. The slide was placed in the thermocycler for 5 minutes to dry. A coverslip was placed on the slide to prepare for imaging with the microscope.

## Results

### Ultrasound

The WSS of the CCA in systole (p = 0.001) and diastole (p = 0.0004) was statistically significant between groups, as shown in Figure 3. During systole, aged 3xTg WSS was increased by 57.7% compared to aged controls (p = 0.0037). Although not statistically significant, young 3xTg WSS was increased by 34.5% compared to young controls; aged 3xTg WSS was increased by 7.33% compared to young 3xTg; and aged control WSS was decreased by 8.44% compared to young controls.

**Figure 3.**
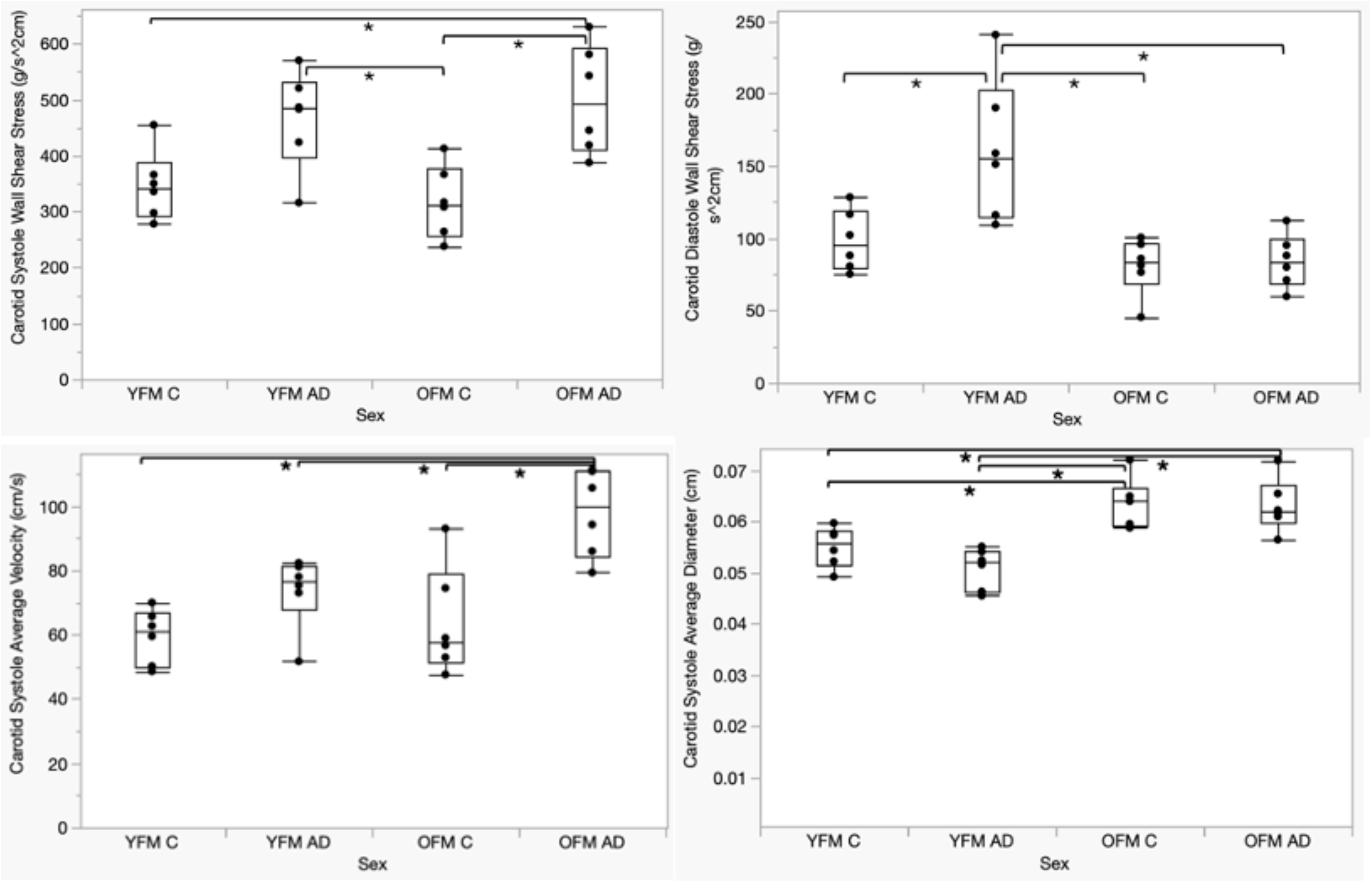
Wall shear stress of the common carotid artery during **(top left)** systole and **(top right)** diastole. Wall shear stress varied across sex and age with significance p<0.05 denoted between groups(*). YFM: young female (12 week), OFM: old female (12 month), C: control, AD: 3xTgAlzheimer model. **(Bottom Left)** Average velocity of the common carotid artery during systole. **(Bottom Right)** Average diameter of the common carotid artery during systole. Diameter and velocity varied with respect to age and AD with significance p<0.05 denoted between groups(*).

During diastole, young 3xTg WSS was increased by 63.5% compared to young controls (p = 0.0083). Although not statistically significant, aged 3xTg WSS was increased by 4.23% compared to aged controls. Aged 3xTg WSS decreased by 47.6% compared to young 3xTg (p=0.0013). Although not statistically significant, aged control WSS decreased by 17.9% compared to young controls.

The PI of the CCA was statistically significant with a value of p = 0.002, as shown in Figure 4. Aged 3xTg PI increased by 26.1% compared to aged controls (p=0.0064). Although not statistically significant, young 3xTg PI decreased by 8.16% compared to young controls. aged controls PI decreased by 4.38% compared to young controls. For aged 3xTg, the PI increased by 31.6% compared to young 3xTg (p=0.0018). Although not statistically significant, aged control PI decreased by 4.05% compared to young controls.

**Figure 4.**
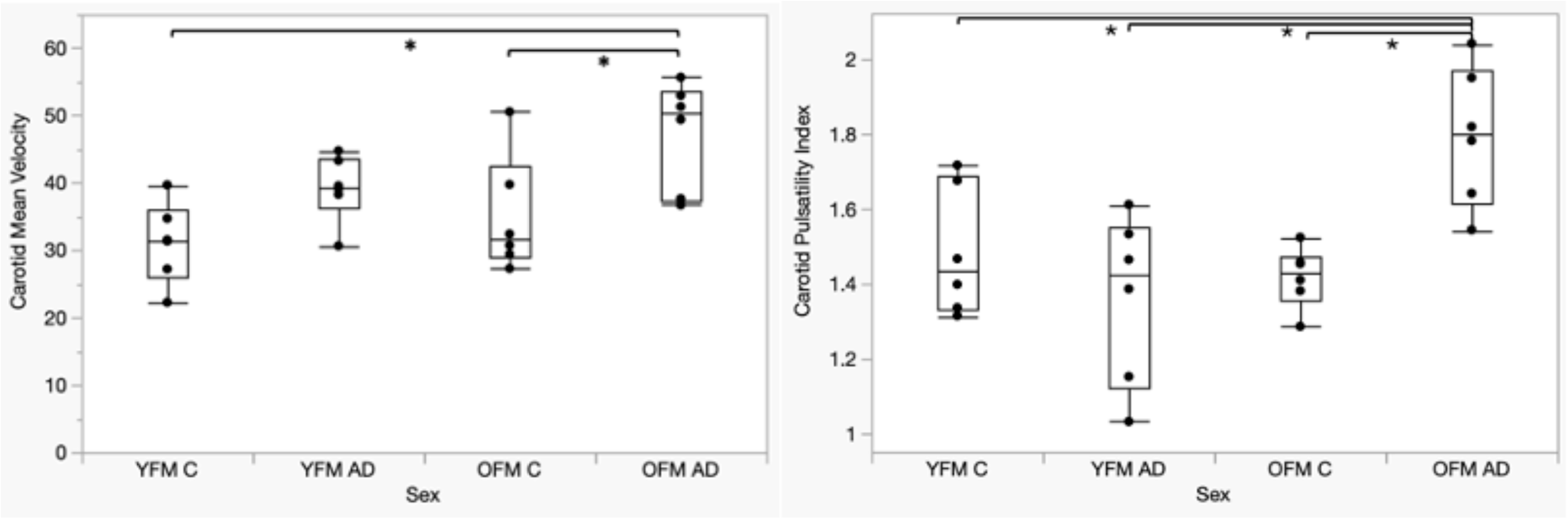
**(Left)** Mean velocity of common carotid artery (CCA) over the cardiac cycle. **(Right)** Pulsatility index of the CCA. Both mean velocity and PI varied across sex and age with significance ofp<0.05 denoted between groups(*)

The CCS values for the carotid arteries and the WSS values for the jugular veins between groups were measured but were not statistically significant. For the left carotid, the CCS was 26.1 ± 12.3 for young controls, 30.2 ± 16.8 for young 3xTg, 16.2 ± 8.85 for old control, and 25.0 ± 12.6 for aged 3xTg. For the right carotid, the CCS was 23.0 ± 7.93 for young controls, 30.5 ± 15.8 for young 3xTg, 22.8 ± 4.91 for aged controls, and 31.0 ± 12.4 for aged 3xTg.

### Mass Spectrometry

MALDI-MSI and LC-MS data of 3xTg and controls were compared. LC-MS results identified several lipids with statistically significant quantities (p<0.05) which can be seen in the PCA and PLS-DA plot in Figure 6. The fold change is the average of all six samples for one group divided by the average of all six samples from a comparable group. The fold change was computed for young AD vs. young control, aged AD vs. aged control, aged control vs. young control, and aged AD vs. young AD. The m/z values from the LC-MS results were compared to the MALDI MSI images to search for similarities.

**Figure 5.**
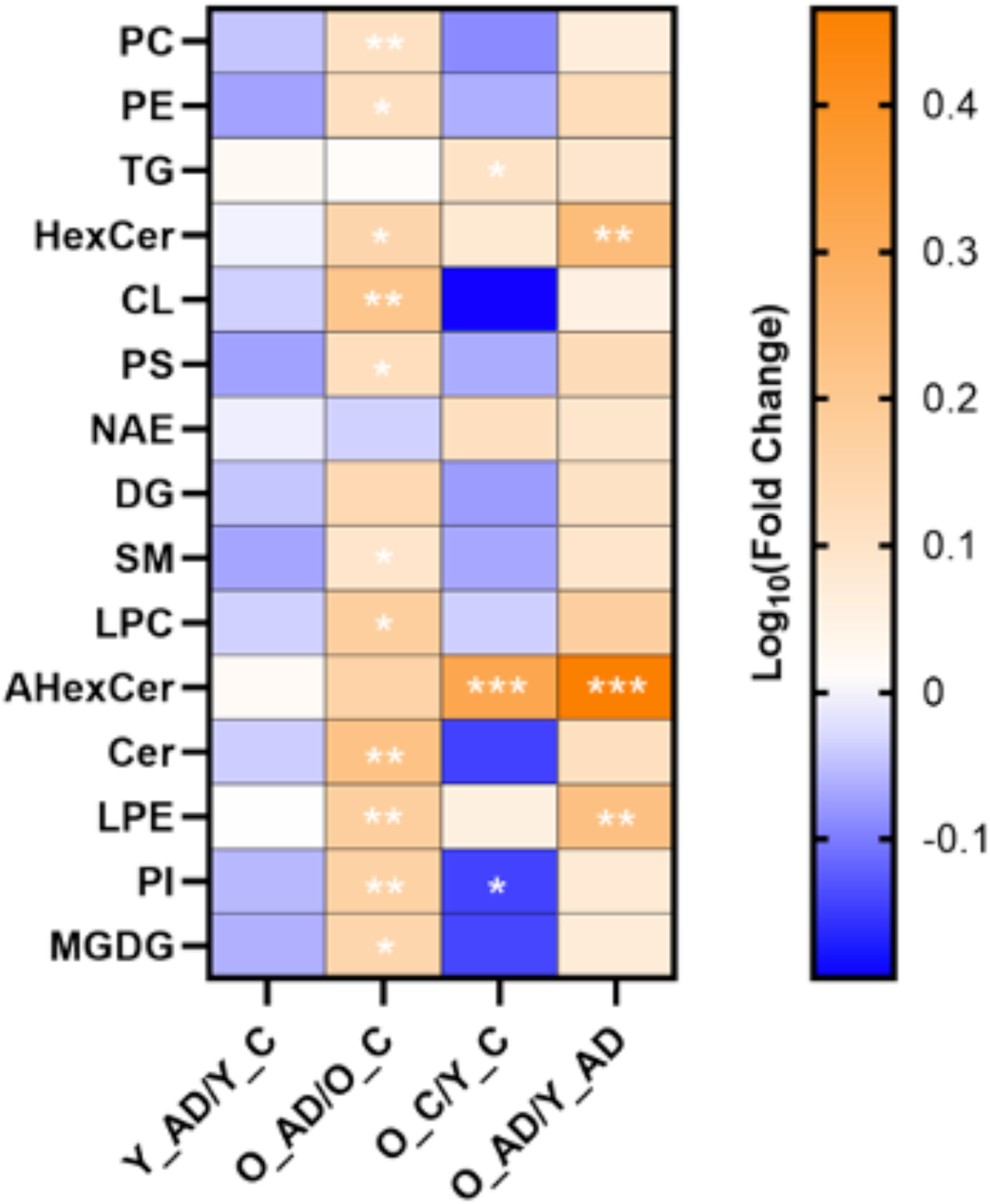
Heatmap of lipid headgroups with 10 or more species detected in LC-MS brain samples. The heatmap provides a visual representation of the fold changes between the four groups listed at the bottom, with significance of p<O.l (*), p<0.05 (**), and p<0.01 (***) denoted between groups.

**Figure 6.**
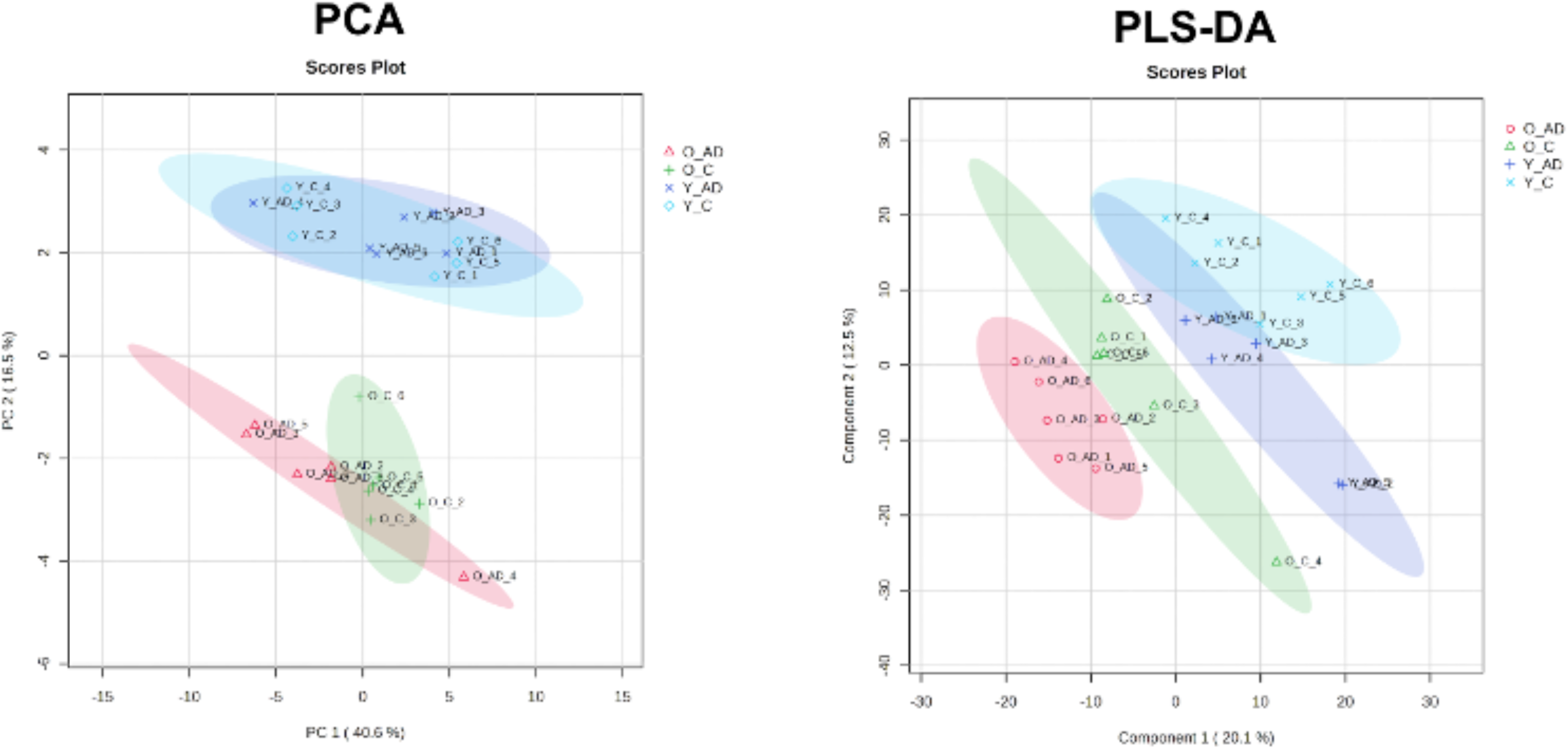
**(Left)** PCA of all four groups showing distinct variation between young and aged groups and between aged control and AD. **Right** PLS-DA of all four groups which emphasizes the variations between all four groups.

The abundance of several lipid headgroups changed significantly with respect to age and AD genotype, as shown in Figure 5. In clinical studies, lysophosphatidylcholines (LPC), phosphatidylethanolamines (PE), and phosphatidylinositol (PI), classes of phospholipids, have been shown to decrease with AD [52][53], [54]. In mouse studies, cardiolipins (CL) have been shown to decrease with AD [53], [55]. In clinical and mouse studies, phosphatidylcholines (PC) were shown to decrease with AD [52]. However, we observed a statistically significant increase in LPCs, PEs, CLs, and PCs in the aged 3xTG mouse model compared to aged controls. In figure 7, PC O-(37:9), a phosphatidylcholine with an m/z of 772.53, is depicted in orange and is primarily located in the hippocampus and outer region of the brain. PC O-(37:9) was found to be more abundant in aged 3xTg compared to aged controls with a fold change of 1.96 and p-value of 0.007. From the MALDI data, PC O-(37:9) was found in all 4 groups and was within the highest average intensities across the entire tissue sample. In figure 8, PE O-42:10, with m/z of 798.54, is pictured in all MALDI samples with the intensity increasing with AD.

**Figure 7.**
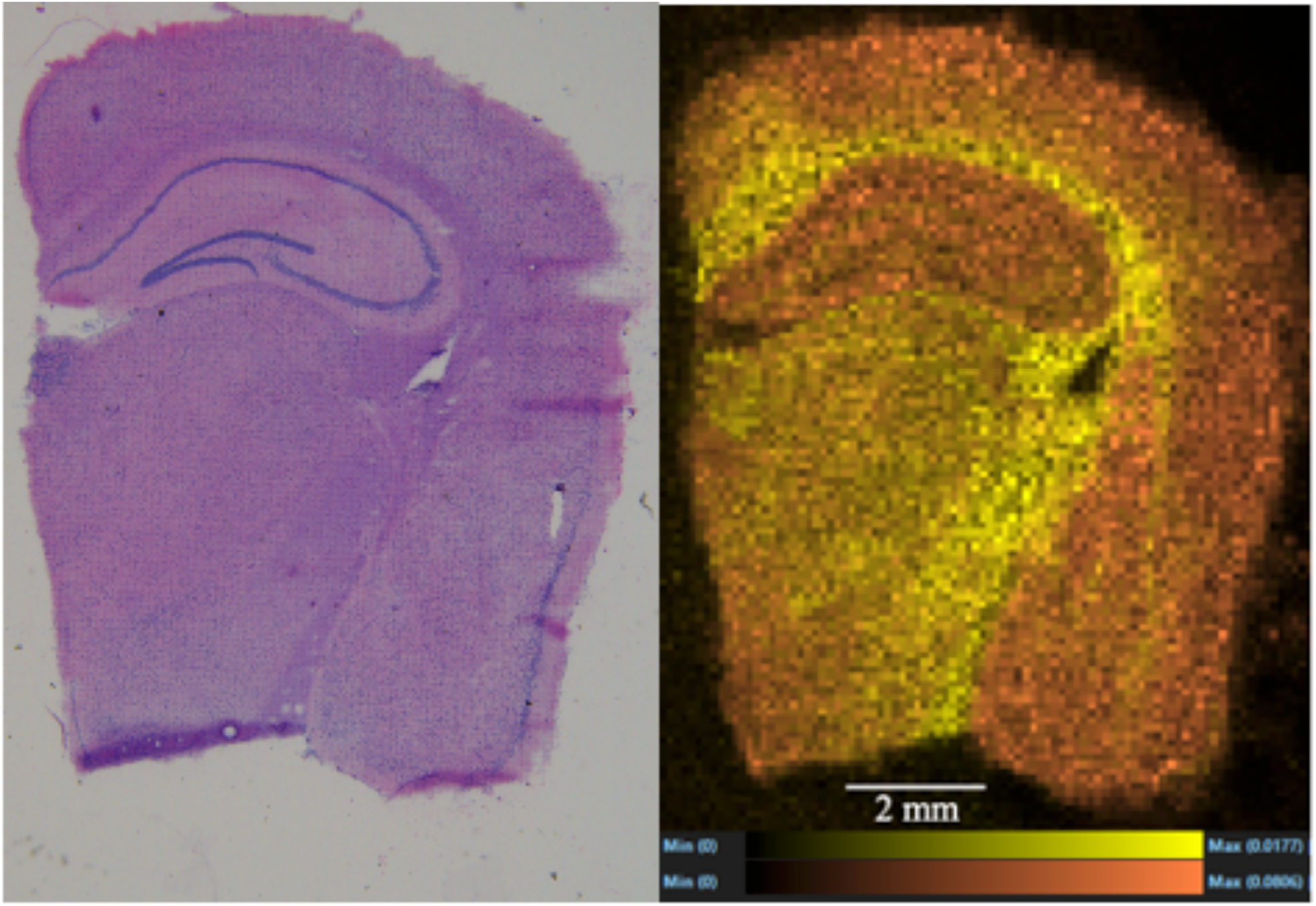
**(Right)** <u>H&E stained</u> microscope image of the right hemisphere of an aged 3xTg brain tissue. **(Left)** Molecular overlay of two different molecules in the same aged 3xTg brain tissue. PC 0-(37:9), a phosphatidylcholine with m/z of 772.53, is depicted in orange and PC 40:4, with m/z of838.62, is depicted in yellow. This image shows the localization of some of the most intense lipids found in the brain tissue.

**Figure 8.**
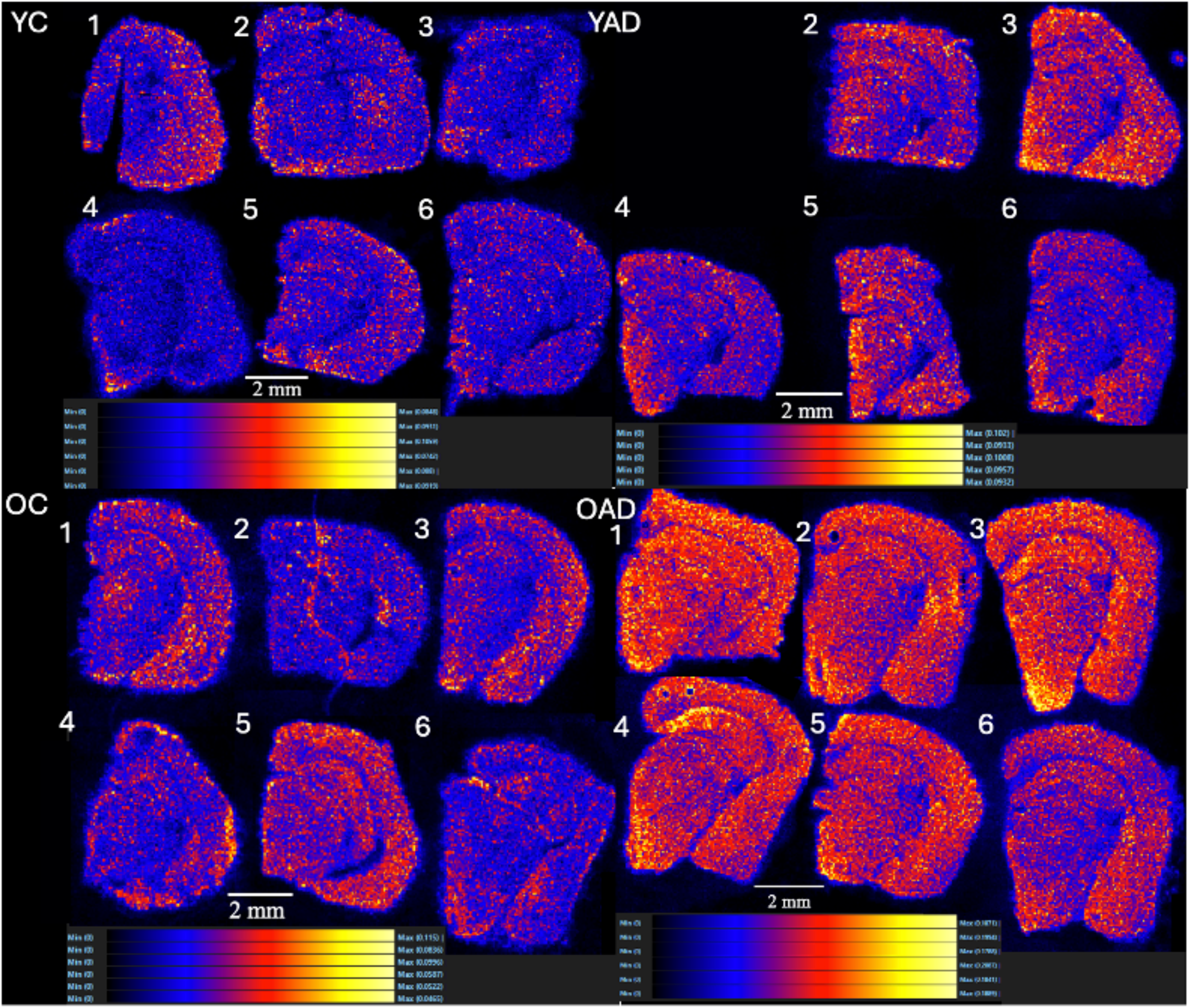
PE 0-42:10, with m/z of798.54, present in the brain tissue ofall 4 groups. Images show intensity and localization of PE 0-42:10 in the brain. **(Top left)** Young control 1-6 **(Top right)** Young 3xTg 2-6 (YAO 1 not pictured due to issue during MALDI MSI data collection. **(Bottom left)** Aged control 1-6 **(Bottom right)** Aged 3xTg 1-6.

In mouse studies, lysophosphatidylcholines (LPC) and phosphatidylserine (PS), classes of phospholipids, have been shown to increase with AD in previous studies[55]. In clinical studies hexosylceramides (HexCer), ceramides (CER) and sphingomyelin (SM), classes of sphingolipids; lysophosphatidylethanolamine (LPE), a class of phospholipids; and onogalactosyl diglycerides (MGDG), a class of glycerolipids, have been shown to increase with AD in previous studies [56][57][53][58], [59] ,which aligns with the statistically significant increase of LPC, HexCer, CER, SM, LPE, and MDG observed in the aged 3xTg group compared to controls.

When comparing the LC-MS data with the MALDI MSI data, there were several lipids found in all six mice within each group in both the MALDI MSI and LC-MS data. LPC 16:0 [M+H]^+^, a lysophosphatidylcholine, was more abundant in aged 3xTg compared to aged controls with a fold change of 1.41. Figure 9 shows the location and intensity of this lipid that was found in all 6 of the aged 3xTg group [60][61]. LPC (18:0), a lysophosphatidylcholine with m/z of 524.37, was more abundant in aged 3xTg compared to young controls with a fold change of 1.57 (p= 0.04) and more abundant in aged 3xTg compared to young 3xTg with a fold change of 1.85 (p= 0.05). LPE 18:1, a lysophosphatidylethanolamine with m/z of 480.31, was more abundant in aged 3xTg compared to young 3xTg with a fold change of 4.68 (p=0.0001), more abundant in aged controls compared to young controls with a fold change of 4.69 (p=7x10^-9^), and more abundant in aged 3xTg compared to aged control with a fold change of 1.47 (p=0.02).

**Figure 9.**
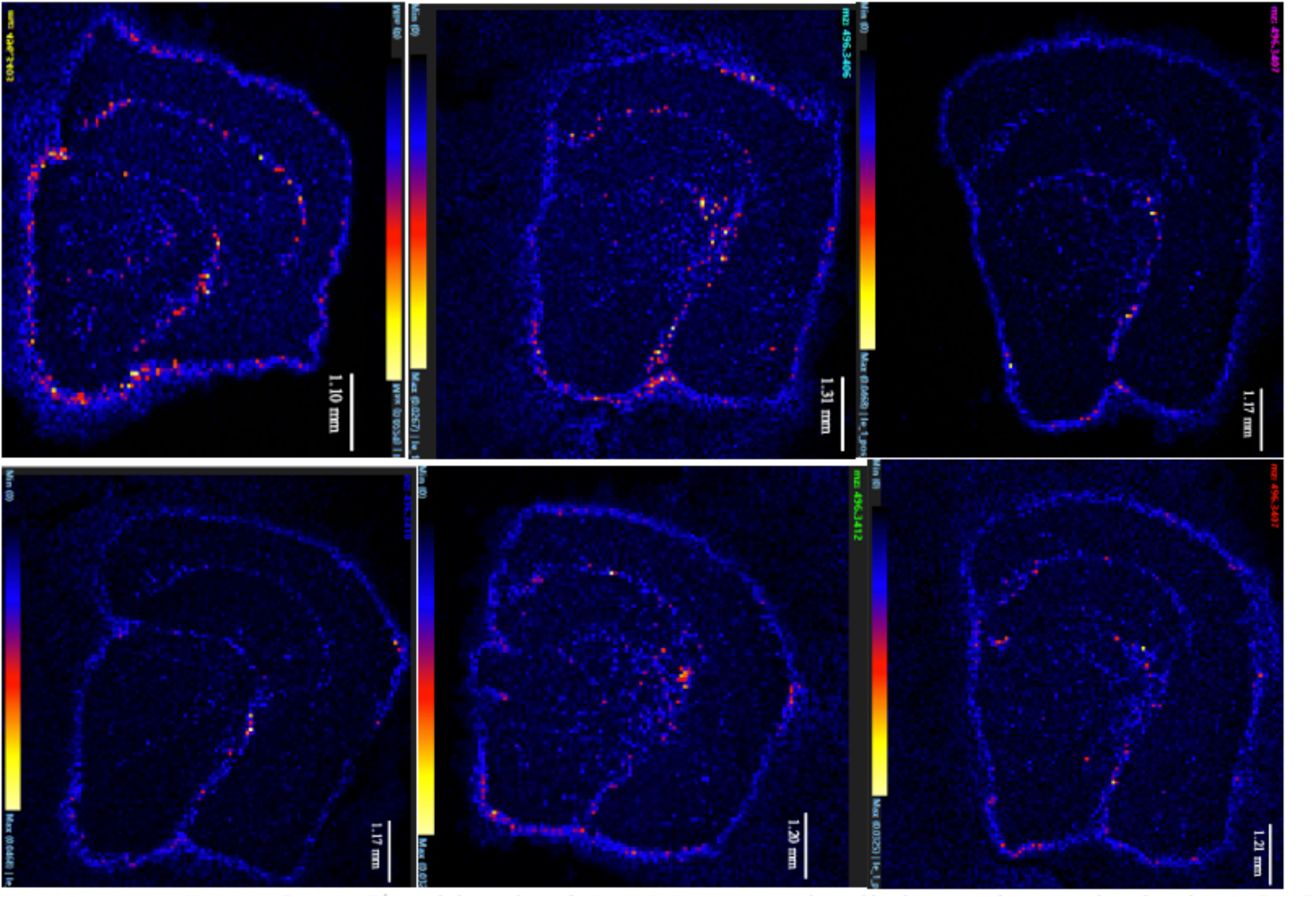
LPC 16:0 <u>[M+HJ+</u>, with m/z of 496.34, present in all six aged 3xTg brain tissue[81]. These images show the localization of this lipid within the right hemisphere of each brain starting with OFM1-OFM3 from left to right in the top row and OFM4-OFM6 from left to right in the bottom row. LPC 16:0 <u>[M+H]+</u>can be seen primarily in the outer region of the hippocampus, a region vital to the detection of Alzheimer’s disease[82].

## Discussion

To study the effect of aging, the two age groups used in this experiment were 12 weeks (young group) and 52 weeks (aged group). The young group corresponds to about a 20-30-year-old human and the aged group corresponds to about a 50–60-year-old human [62]. In the 3xTg mouse model, signs of cognitive decline have been detected as early as 3-5 months of age with the first signs of associate learning deficits [63]. At around 6 months of age, impairments in spatial working memory begin followed by deficits in recognition memory at around 9-11 months. At around 12 months, reference memory impairment is detected [63].

### Ultrasound

Age-related arterial stiffness can cause increased flow pulsatility in the microvasculature, potentially leading to cognitive decline and AD [64]. The microvasculature that consists of thin-walled arteries, arterioles, and capillaries is not designed to withstand high velocities, increasing the risk of damage. Since the brain is highly susceptible to fluctuations in blood pressure, the increase in velocity caused by vascular stiffening can have a significant effect, possibly leading to damage of vital regions in the brain [5]. Over time, the constant high velocity and pressure in the cerebral arteries could lead to endothelial dysfunction and breakdown of the blood brain barrier initiating cognitive decline indicative of AD [65] [66].

To deepen our understanding of the relationship between vascular stiffening and AD, the carotid and jugular were imaged via ultrasound while the brain was imaged using MALDI MSI. CCS has been shown to decrease with signs of vascular stiffening, however, there was no significant change from our ultrasound results[67]. The observed trend seen in CCS was a decrease with age but an increase with AD. The lack of statistically significant results may be attributed to the group’s insufficient maturation at 12 weeks. A previous study observed significant changes seen in CCS at about 18 weeks of age [68]. In previous studies, WSS has been shown to decrease with age causing endothelial dysfunction and arterial stiffness[69], [70]. During systole, our ultrasound results show a significant increase in WSS with AD in the aged group. Without the confounding influence of aging, the change in WSS was not significant between the control and AD groups. During diastole, our results show a significant decrease in WSS with aging and AD. Without the presence of AD, WSS did not significantly change with age. As seen in figure 5, the increase in WSS of the carotid during systole between aged controls and aged 3xTg was driven by a significant increase in average velocity during systole. The average velocity between young and aged control remained relatively unchanged. This leads us back to our hypothesis that vascular stiffening, which leads to an increase in velocity in the large arteries, may contribute to AD. Since the large arteries lead directly to the microvasculature surrounding the brain, any fluctuation in blood flow can greatly impact the microvasculature and, therefore, lead to a disruption in the blood-brain barrier.

Pulsitility index has been shown to increase with age and AD [71], [72]. Higher PI has been associated with increased vascular resistance and hypertension [73]. Our ultrasound results show a significant increase in PI with AD in the aged group, but no significant change with AD in the young group. PI increased significantly with age in the AD groups, but there was no significant change with age in the control groups. Our results indicate a significant increase in PI with the simultaneous presence of AD and aging. The increase in PI seen in the aged AD group compared to aged controls indicates the potential presence of vascular dysfunction in the AD group.

### Mass Spectrometry

Since lipids are the mediators that manage many immune responses, they are vital when studying a disease such as AD, which may be associated with divergent immune responses [24]. Lipids indicative of AD tend to be localized in different regions of the brain, which can be visualized using MALDI MSI data [13]. LPC 16:0 was elevated in the aged 3xTg group compared to the aged control. In figure 10, LPC 16:0 is shown in all six aged 3xTg brain tissue and is primarily localized in the olfactory tract, which is a region that plays a crucial role in our sense of smell [74]. Loss of smell has been shown to have a positive correlation with signs of cognitive impairment indicative of AD [75]. LPCs play an important role in inflammatory response and the localization of LPC 16:0 in the olfactory tract may contribute to inflammation affecting our sense of smell [76][77]. LPC 18:0 was found in all six aged 3xTg in both the LC-MS and MALDI MSI data and has been shown to increase with signs of plaque formation in atherosclerosis and AD [78], [79]. LPE 18:1 was found in all six aged 3xTg and has been associated with a quicker progression from mild cognitive impairment to AD [59].

PCs were elevated in the aged AD group compared to aged controls. PCs plays a role in cell signaling and are the primary components of neuronal cell membranes [52]. Table 3 summarizes the abundances of specific lipids identified in our dataset with respect to AD, emphasizing their consistency or inconsistency with findings from previous studies. As seen in Table 3, there were several lipids that aligned with the results of previous studies both in human and rodent models. Most of the inconsistent findings are most likely due to variation in the model studied, highlighting the need to have baseline molecular characterization of AD models before studying novel treatments or comparing to humans. Table 4 summarizes the abundance of specific lipids identified in our dataset with respect to age and whether or results align with the results of previous studies. Our findings indicate that the abundance and location of certain lipids significantly varied with age and AD.

A few limitations were present in this study. First, using mouse models to study a disease present in humans such as AD creates some limitations since they are not biologically identical to humans. Second, the 3xTg mouse model should not be used for sex comparisons in AD because the males do not develop the phenotypic traits of AD [25], [28], [29]. Third, the ultrasound measurements were performed with the animals under anesthesia using isoflurane, which can potentially reduce signs of stiffening [80].

In conclusion, our work emphasizes the differences in vascular biomechanics and brain biochemistry seen with aging and AD. Our ultrasound results indicate significant changes in WSS, PI, velocity, and diameter of the carotid arteries with the presence of AD. From our LC-MS results, several lipids were significantly changed with the presence of aging and AD. Our MALDI MSI results were able to identify the spatial distribution and intensities of lipids across the brain tissue. Future studies are needed to investigate how sex differences play a role in the relationship between the vasculature and brain biochemistry. Understanding how sex differences affect the onset of AD could explain the increased prevalence of AD in women compared to men.

| Lipid | M/z | Group | Consistent | Model | Inconsistent | Model |
| --- | --- | --- | --- | --- | --- | --- |
| SHexCer 42:1;O3 | 908.648 | OAD > OC |  |  | OAD < OC<br>(Zhang 2024) | Mouse |
| PC 32:0 | 756.55 | OAD < OC | OAD < OC<br>(Sabogal-Guáqueta, 2018)<br>OAD < OC<br>(Whiley, 2014)<br>OAD < OC<br>(Zhang, 2021) | Rats<br>Clinical<br>Mouse |  |  |
| SM 36:1 | 753.58 | OAD < OC |  |  | OAD > OC.<br>(Koal, 2015) | Clinical |
| PC O 36:3 | 770.607 | OAD > OC | OAD > OC<br>(Puris, 2022) | Mouse |  |  |
| TG 56:8 | 920.773 | OAD < OC | MCI & AD < Control<br>(Bernath, 2020) | Clinical |  |  |
| Cer 22:0 | 370.332 | OAD < OC |  |  | MCI > Control<br>(Wackerlig, 2020) | Clinical |
| PC 38:1 | 816.648 | OAD > OC | MCI > Control<br>(Abdullah, 2012) | Mouse |  |  |
| PC 40:4 | 838.63 | OAD > OC | MCI > Control<br>(Abdullah, 2012)<br>AD > Control<br>(Proitsi, 2021) | Mouse |  |  |
| PC 38:6 | 806.568 | OAD > OC |  |  | OAD < OC<br>(Whiley, 2014) | Clinical |
| PC 40:6 | 834.599 | OAD > OC<br>OC < YC |  |  | OAD < OC<br>(Whiley, 2014) | Clinical |
| Cer 44:2 | 708.651 | OAD > OC |  |  | MCI > Control<br>(Marković, 2024) | Clinical |
| PE O-42:10 | 798.54 | OAD > OC<br>OC < YC | AD > Control<br>(Fernandes, 2024) | Cells |  |  |

| Lipid | m/z | Group | Consistent | Model | Inconsistent | Model |
| --- | --- | --- | --- | --- | --- | --- |
| Cer 22:0 O2 | 404.33 | OAD>YAD | Old>Young<br>(Crivelli 2024) | 5xFAD |  |  |
| Cer 22:3 O3 | 370.332 | OAD>YAD<br>OC>YC | Old>Young<br>(Crivelli 2024) | 5xFAD |  |  |
| Cer 20:0 | 344.315 | OAD>YAD | Old>Young<br>(Crivelli 2024) | 5xFAD |  |  |
| PC O 36:3 | 770.607 | OAD>YAD | Old>Young<br>(Panda, 2024) | C57BL/6 |  |  |
| DG 38:4 | 662.573 | OAD>YAD | Old>Young<br>(Panda, 2024) | C57BL/6 |  |  |
| PC 38:3 | 814.55 | OC<YC |  |  | Old>Young<br>(Panda, 2024) | C57BL/6 |
| HexCer 40:1 | 800.661 | OC>YC<br>OAD>YAD | Old>Young<br>(Panda, 2024) | C57BL/6 |  |  |

## Notes

### Competing Interest Statement

The authors have declared no competing interest.

